# Survivin Promotes the Formation of a Microtubule-Based Glycolytic Hub

**DOI:** 10.64898/2026.08.28.747899

**Authors:** Jacob Neumann, Wen-Hsuan Chang, Sarah E. Ackermann, Matthew R. Zanotelli, Tomer Markovich, Runying Yang, Jolie R. Lefkowitz, Shun Enomoto, Henry H. Le, Min-Ting Lee, Kirsten L. Bryant, Richard A. Cerione, Marc A. Antonyak

**Affiliations:** Department of Chemistry and Chemical Biology, Cornell University, Ithaca, New York, USA; Lineberger Comprehensive Cancer Center, University of North Carolina at Chapel Hill, Chapel Hill, North Carolina, USA; Department of Pharmacology, University of North Carolina at Chapel Hill, Chapel Hill, North Carolina, USA; Department of Biomedical and Translational Sciences, Cornell University, Ithaca, New York 14853, USA; Division of Nutritional Sciences, Cornell University, Ithaca, NY, USA

**Keywords:** Survivin, Glycolysis, Microtubules

## Abstract

KRAS is one of the most frequently mutated oncoproteins in cancer. Its ability to induce malignant transformation relies on metabolic reprogramming that causes cells to become dependent on aerobic glycolysis as a primary source of energy and for generating biological building blocks. Thus far, the signaling mechanism used by oncogenic KRAS to promote these changes in cancer cell metabolism has not been fully elucidated. However, through studies in pancreatic ductal adenocarcinoma (PDAC) cell lines and patient-derived organoids, we now demonstrate how oncogenic KRAS triggers an increase in glycolytic activity and identify Survivin as a newly discovered and critical KRAS-signaling partner essential for promoting these metabolic changes. We show that oncogenic KRAS potently upregulates the expression of Survivin in PDAC cells and patient-derived organoids undergoing increased glycolysis, whereas depleting Survivin expression inhibits their glycolytic activity and growth. Through a combination of cellular, biochemical, and imaging approaches, we further show that Survivin promotes the formation of unique microtubule-based structures that resemble invadosome rosettes, allowing for the recruitment of the glycolytic enzymes triose phosphate isomerase (TPI) and glyceraldehyde-3-phosphate dehydrogenase (GAPDH) to these super-structures which drives the increases in glycolysis. These findings demonstrate that by directing the assembly of a microtubule-based complex of metabolic enzymes, Survivin serves as a vital link in a KRAS signaling pathway responsible for promoting the metabolic changes necessary for the accelerated growth of PDAC cells, and thus potentially highlight new therapeutic strategies for treating KRAS-dependent cancers.

**Significance Statement:** Oncogenic forms of the signaling protein KRAS promote the rapid growth of pancreatic cancer cells by rewiring their cellular metabolism such that they become dependent on aerobic glycolysis. However, how this metabolic adaptation is achieved has remained a critical open question in the field. In this study, we now show that there is a tight signaling connection between oncogenic KRAS and increases in the expression of the cell survival protein/cancer marker Survivin in pancreatic cancer cell lines and patient-derived organoids, with the upregulation of Survivin being essential for promoting aerobic glycolysis. The mechanism underlying this effect involves the formation of a unique Survivin-microtubule-based metabolic hub that enables the recruitment of key enzymes responsible for increasing glycolytic activity and cell growth.

## Introduction

Approximately 30% of all cancers have mutations in KRAS (for <u>K</u>irsten <u>R</u>at Sarcoma), and in specific types of cancer, the incidence of KRAS mutations is significantly higher [1]. For example, KRAS is mutated in more than 90% of all PDAC tumors, a particularly aggressive and deadly form of cancer that continues to be very difficult to treat [2–4]. These mutated forms of KRAS are constitutively active and result in the increased activation of key signaling effectors, including extracellular signal-regulated kinase (ERK) and phosphoinositide 3-kinase (PI3 kinase) [2–5], that alter the metabolism and enhance the growth of cancer cells resulting in malignant transformation [6,7]. One of the most prevalent changes that KRAS-transformed cancer cells undergo is a reprogramming of their glucose metabolism from oxidative phosphorylation to aerobic glycolysis, a process referred to as the Warburg effect [8]. Although it is still not fully understood how mutant forms of KRAS increase glycolytic activity, it has been well established that cancer cells adopt this change in their metabolic program to generate nucleic acids, proteins, and membranes which are necessary to sustain their rapid growth [9–10]. Attempts to target KRAS, one of its downstream signaling effectors, or even glycolysis, as a treatment for KRAS-dependent cancers like PDAC have been aggressively pursued but thus far have yielded only limited success in the clinics [11–13]. One exception is the new pan-RAS inhibitor Daraxonrasib, which was recently shown in clinical trials to increase the medium survival time of PDAC patients from ∼7 months to ∼13 months [14].

There is a tight signaling connection between oncogenic KRAS and the expression of Survivin, also referred to as baculoviral inhibitor of apoptosis repeat-containing 5 (BIRC5), a member of the inhibitors of apoptosis (IAP) family [15–18]. While there is little or no detectable expression of Survivin in most adult tissues and differentiated cells, it is highly expressed in several types of tumors and cancer cell lines [17–19]. Indeed, tumor sections obtained from PDAC patients and analyzed for Survivin expression by immunohistochemistry showed that 88% of the tumors exhibited a striking increase in Survivin expression compared to adjacent normal tissue [19]. We and others have shown that the high levels of Survivin expression in PDAC cells are lost when inhibiting the activity of oncogenic KRAS mutants, or by targeting one of their major effectors, ERK [17, 18]. Moreover, depleting cancer cells of Survivin upon treatment with either a small molecule that blocks its transcription, or Survivin-targeting siRNAs and shRNAs, potently inhibits their growth, ability to survive stresses, as well as their migration and invasion [17, 20–24]. These findings, combined with the marked increases in Survivin expression being used to diagnose and monitor disease progression, as well as predict therapy responses in cancer patients [25], all highlight Survivin as having a key role in cancer progression.

Despite its extensive use as a cancer marker, how Survivin contributes to malignant transformation has been difficult to determine, given that it lacks enzymatic activity or any discernible scaffolding function [15, 16]. Perhaps the best-described mechanism for how Survivin promotes the growth of cells involves its ability to associate with microtubules that make up mitotic spindles that form during mitosis [22]. This interaction has been proposed not only to maintain the structural integrity of mitotic spindles, but by serving as a component of the chromosome passenger complex (CPC), to tether chromosomes to the mitotic spindles to ensure their proper segregation into daughter cells [15,16]. There also have been suggestions that Survivin stimulates cell proliferation by blocking the actions of a negative regulator of the cell cycle, p16INK4a [23], and preventing the degradation of the transcription factor and proto-oncogene c-MYC [17].

Here, we set out to determine the consequences of oncogenic KRAS-induced Survivin expression, which led us to discover a previously unknown but essential role for Survivin in malignant transformation. We show that depleting PDAC cell lines and patient-derived organoids of Survivin significantly reduces their glycolytic activity, thus inhibiting their growth. We demonstrate that this is due to the formation of a unique Survivin-microtubule-based complex, which enables key glycolytic enzymes including TPI and GAPDH to more efficiently localize to microtubules which gives rise to a significant increase in glycolytic activity. Thus, taken together, these findings highlight how Survivin plays a previously unappreciated but critical role in the ability of oncogenic KRAS to increase glycolytic activity which is essential for satisfying the metabolic requirements necessary for the rapid growth of cancer cells.

## Results

### Identifying an essential role for Survivin in promoting aerobic glycolysis

To investigate the consequences of the upregulated expression of Survivin induced by oncogenic KRAS, we used PDAC cells given that they are derived from a KRAS-driven form of cancer [2–4]. Fig. 1a shows that when MIA PaCa-2 cells were treated with AMG510 (Sotorasib), an inhibitor that targets the KRAS G12C mutant [26], there was a striking reduction in the ability of oncogenic KRAS to activate effectors, including ERK (*compare lanes 1 and 2*). The cells were then assayed for glycolytic activity by determining the amount of lactate, the final product of aerobic glycolysis [8,27], that is secreted into their medium. Compared to cells treated with DMSO, which serves as a vehicle control, the AMG510-treated cells showed an ∼50% reduction in glycolytic activity (Fig. 1b). Similar results were obtained when ERK activity was blocked using the small molecule ERK1/2 inhibitor SCH772984 (Figs. 1a, *compare lanes 1 and 3,* and 1b). These treatments also potently inhibited the growth of the cells (Fig. 1c).

**Figure 1.**
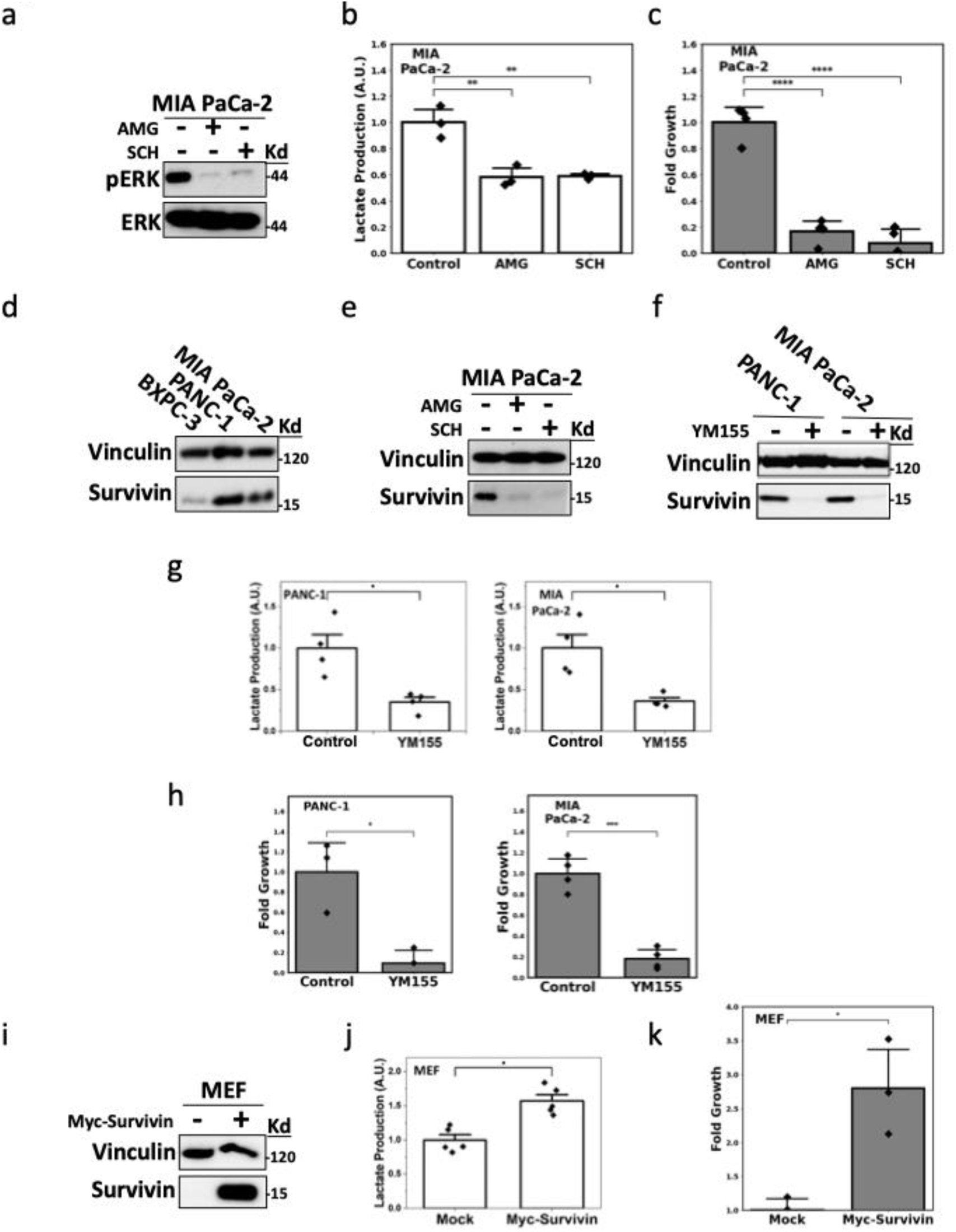
**a.** Western blot showing phosphorylated and total ERK levels (pERK and ERK) in MIA PaCa-2 cells treated without (-; DMSO only) or with (+) 1 µM AMG510 (AMG) or 1 µM SCH772984 (SCH). **b and c.** Lactic acid (b) and growth assays (c) were performed on MIA PaCa-2 cells treated as described in (a). **d-f.** Western blots showing Survivin expression in (d) different PDAC cell lines, (e) MIA PaCa-2 cells treated as described in (a), and (f) PDAC cell lines treated without (-; DMSO only) or with (+) 100 nM YM155. **g and h.** Lactic acid (g) and growth assays (h) were performed on PANC-1 and MIA PaCa-2 cells treated without (Control; DMSO only) or with 100 nM YM155. **i**. Western blot showing Survivin expression levels in mouse embryonic fibroblasts (MEFs) transfected with either an empty vector (-) or a vector encoding Myc-tagged Survivin (+). **j and k.** Lactic acid (j) and growth assays (k) were performed on MEFs treated as described in (i). All experiments shown were performed at least three independent times, and vinculin and ERK were used as loading controls for the Western blots. The data in **b, c, g, h, j**, and **k** are presented as the mean, with p-values denoted as follows: * < 0.05, ** < 0.01, *** < 0.001, **** < 0.0001. Error bars indicate one standard deviation (SD).

Survivin is highly expressed at the protein level in two oncogenic KRAS-driven PDAC cell lines [17], PANC-1 and MIA PaCa-2 (Fig. 1d), while an examination of the TCGA database shows that the transcript encoding Survivin, referred to as BIRC5, is also markedly upregulated in the vast majority of PDAC tumor samples (*Supporting Information (SI) Appendix*, Fig. S1a). However, an exception is the BxPC3 cell line, which is one of the few PDAC cell lines that contains a wildtype form of KRAS rather than an oncogenic variant [28], and consequently expresses Survivin at much lower levels, compared to PANC-1 and MIA PaCa-2 cells (Fig. 1d, *compare lanes 1 to 2 and 3*). In MIA PaCa-2 cells, whose transformation is driven by the oncogenic KRAS G12C mutant, treatment with either AMG510 or SCH772984 caused a dramatic reduction in Survivin expression (Fig. 1e). Similar results occurred when PANC-1 cells were treated with SCH772984 (*SI Appendix,* Fig. S1b). It was the ability of oncogenic KRAS signaling to ERK to induce both aerobic glycolysis and Survivin expression, that led us to examine whether Survivin contributes to the metabolic reprogramming of cancer cells. PANC-1 and MIA PaCa-2 cells were depleted of Survivin expression by treatment with either YM155, which blocks Survivin transcription [24], or a Survivin-targeting shRNA (Fig. 1f and *SI Appendix,* Fig. S1c), and then assayed for their glycolytic activity. Similar to the results obtained when inhibiting oncogenic KRAS and ERK activity, PANC-1 and MIA PaCa-2 cells depleted of Survivin showed a 50-60% decrease in glycolytic activity (Fig. 1g and *SI Appendix,* Fig. S1d), accompanied by a striking growth inhibition (Fig. 1h and *SI Appendix,* Fig. S1e). The effects of Survivin on glycolysis and cell growth were not limited to PDAC cell lines, as inhibiting the high levels of Survivin expression in triple-negative MDA-MB-231 breast cancer cells, which express an oncogenic KRAS G13D mutation, with YM155 (*SI Appendix,* Fig. S1f) also resulted in a strong reduction in glycolytic activity and cell growth (*SI Appendix,* Figs S1g and S1h).

To further demonstrate the important role played by Survivin in promoting glycolysis, MYC-tagged Survivin was transiently transfected into a non-transformed cell type, mouse embryonic fibroblasts (MEFs) that typically show very little detectable expression of endogenous Survivin (Fig. 1i). Cells ectopically expressing Survivin showed significantly higher levels of glycolytic activity as read-out by lactate production compared to mock-transfected MEFs (Fig. 1j). The cells expressing Survivin also exhibited a marked increase in their growth compared to the mock-transfected MEFs (Fig. 1k). We next determined whether Survivin plays an important role in promoting aerobic glycolysis and stimulating cell proliferation in more clinically relevant patient-derived PDAC organoids [29]. Four different organoids were examined that were previously shown to express oncogenic KRAS mutants. Specifically, PT6 and PT8 expressed the KRAS G12V mutant, while hT105 and hM1A expressed the KRAS G12D mutant. Sequencing of tumor protein 53 (p53), cyclin-dependent kinase inhibitor 2A (CHKN2A), and mothers against decapentaplegic homolog 4 (SMAD4), was also performed and their mutational status within the organoids is listed in Fig. 2a. Each organoid exhibited Survivin expression (Fig. 2b), albeit at varying amounts, with the relative effectiveness of YM155 treatment lowering their levels (Fig. 2c) being accompanied by a corresponding inhibition of their growth (Fig. 2d). We then assessed how inhibiting Survivin expression affected glycolytic activity in the PT6 organoid which produced considerable amounts of lactate (Fig. 2e), determined by ^1^H-NMR analysis of their conditioned medium as previously described [29] and found that depleting Survivin expression reduced their lactate levels by ∼50% (Fig. 2e).

**Figure 2.**
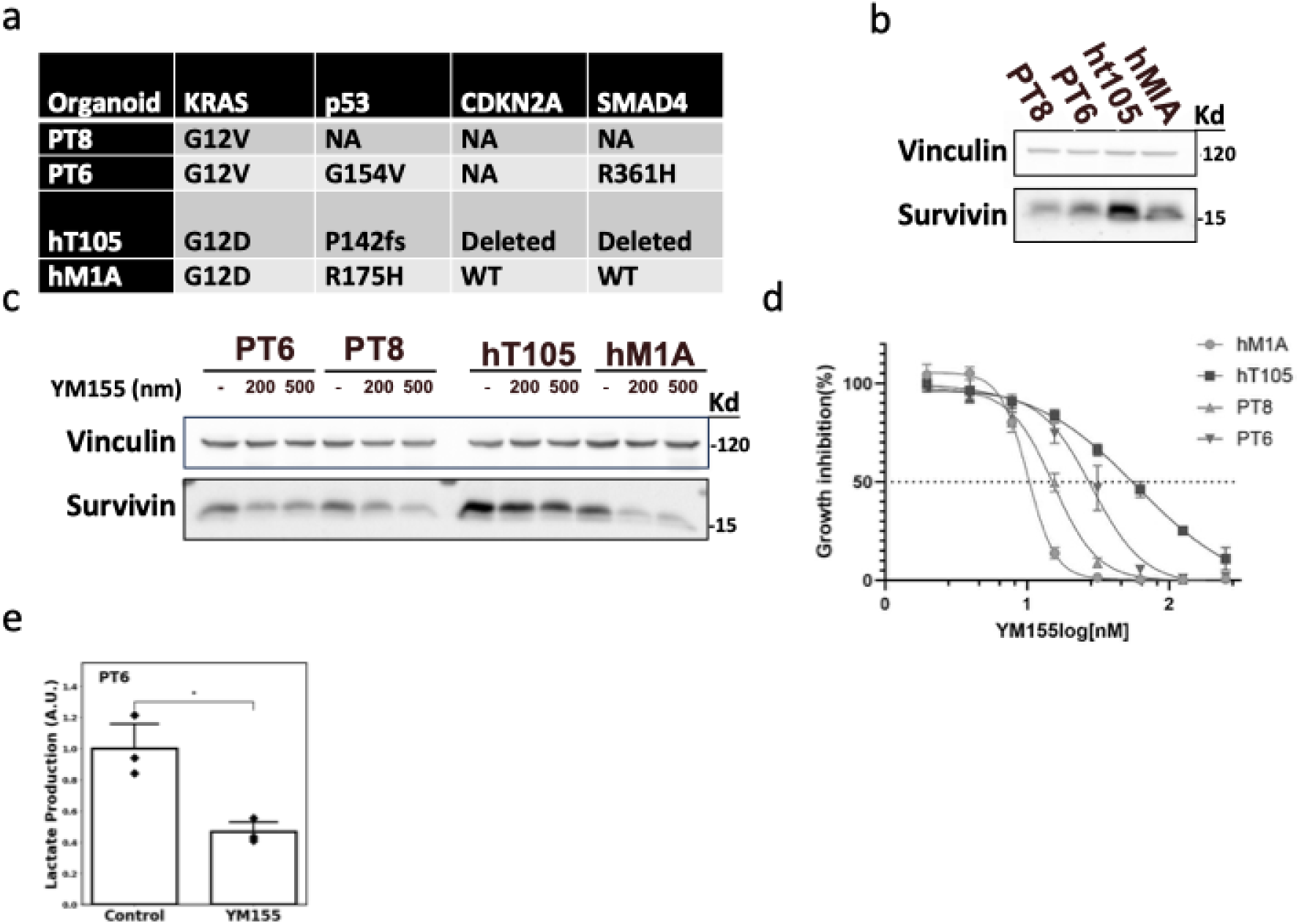
**a.** List showing the names of four different patient-derived organoids (Organoid) and their mutational status of KRAS, p53, CDKN2A, and SMAD; Not available (NA), wildtype (WT), frame shift (fs), and specific point mutants are indicated. **b and c.** Western blots showing Survivin expression levels in the organoids (b) or after the organoids were treated with the indicated concentrations of YM155 (c). **d.** Growth assays were performed on the organoids treated with increasing concentrations of YM155. **e**. ¹H nuclear magnetic resonance (¹H NMR) spectroscopy was performed on the conditioned medium collected from the PT6 organoid to determine its levels of lactate when treated without (Control; DMSO only) or with 100 nm YM155 for 10 hours. All experiments shown were performed at least three independent times, and vinculin was used as a loading control for the Western blots. The data in **d** and **e** are presented as the mean, with p-values denoted as follows: * < 0.05. Error bars indicate one standard deviation (SD).

### Survivin promotes TPI activity

Since glycolysis was decreased by ∼50% in PDAC cells and patient-derived organoids upon their treatment with either the oncogenic KRAS inhibitor AMG510, the ERK inhibitor SCH772984, the Survivin transcriptional inhibitor YM155, or with Survivin-targeting shRNA, we determined whether this degree of reduction in metabolic activity was responsible for the accompanying inhibition that occurred in cell growth. MIA PaCa-2 cells were treated with different concentrations of 2-deoxyglucose (2-DG), a non-hydrolysable glucose analog that blocks the first step in glycolysis [30], to determine the amount required to inhibit glycolytic activity to a comparable extent as observed when depleting cells of Survivin (i.e. a reduction of ∼50%). Fig. 3a shows that 2-DG at a concentration of 5.0 mM was sufficient to reduce lactate production by 40-50% and we found that this concentration of 2-DG in cell growth assays potently inhibited the growth of MIA PaCa-2 cells (Fig. 3b).

**Figure 3.**
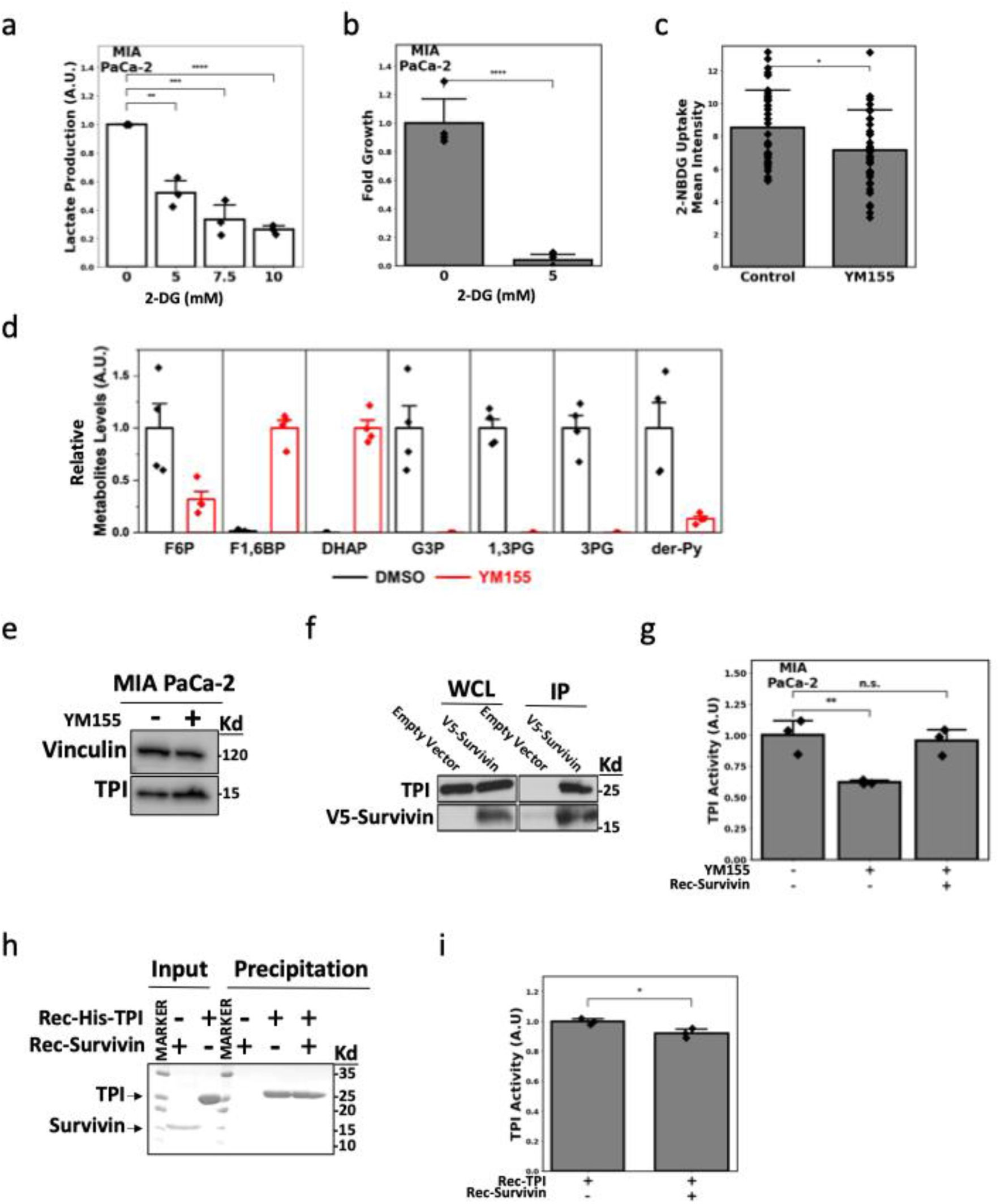
**a and b.** Lactic acid (a) and cell growth (b) assays were performed on MIA PaCa-2 cells treated with the indicated concentrations of 2-DG**. c**. 2-NBDG uptake assays were performed on MIA PaCa-2 cells treated without (Control; DMSO only) or with 100 nM YM155 for 10 hours. **d.** Relative metabolite levels of the indicated glycolytic intermediates were determined using LC-MS/MS in MIA PaCa-2 cells treated without (black lined bars; DMSO only) or with 100 nM YM155 (red lined bars). **e.** Western blot showing TPI expression levels in MIA PaCa-2 cells treated without (-; DMSO only) or with (+) 100 nM YM155. **f.** Western blot showing TPI and V5-tagged Survivin expression levels in the whole-cell lysates (WCL), and immunoprecipitations performed on these lysates using a V5 antibody (IP), from PDAC cells transfected with either an empty vector or a vector encoding V5-tagged Survivin. **g**. TPI activity assays were performed on MIA PaCa-2 cells treated without (-; DMSO only) or with (+) 100 nM YM155. An additional condition was included in this experiment where a purified recombinant form of Survivin (Rec-Survivin) was added to lysates of YM155-treated cells. **h.** Protein precipitation experiments using nickel beads were performed on samples containing the indicated combinations of recombinant forms of 6x His-tagged TPI (Rec-His-TPI) and Survivin (Rec-Survivin). The proteins used are shown (*Input*), as are the results of the precipitation assay (*Precipitation*). **i.** TPI activity assays were performed on samples containing recombinant TPI alone (Rec-His-TPI) or combined with recombinant Survivin (Rec-Survivin). All experiments shown were performed at least three independent times. The data in **a**, **b**, **c**, **g**, and **i** are presented as the mean, with p-values denoted as follows: n.s. not significant, * < 0.05, ** < 0.01, *** < 0.001, **** < 0.0001. Error bars indicate one standard deviation (SD).

Having established that the increase in glycolytic activity caused by Survivin is critical for the growth of PDAC cells, we then set out to elucidate the mechanism responsible for mediating this effect. We first examined whether glucose uptake in MIA PaCa-2 cells was decreased by the depletion of Survivin. However, Fig. 3c shows that uptake of the fluorescent glucose analog 2-NBDG by cells was only minimally affected by treatment with YM155 and thus cannot account for the effects on glycolysis that occur upon the depletion of Survivin expression.

We next performed a metabolomic analyses of MIA PaCa-2 cells that were treated with either DMSO (i.e., the vehicle control) or YM155 to block Survivin expression. Distinct differences were detected in the levels of several metabolites generated in the glycolytic pathway when comparing the two treatment groups (Fig. 3d). Cells expressing Survivin (i.e., cells treated with the vehicle control DMSO; *black bars*) showed high levels of fructose-6-phosphate (F6P), low levels of fructose 1,6-bisphosphate (F1,6BP) and dihydroxyacetone phosphate (DHAP), and relatively high amounts of several subsequent key intermediates formed in the glycolytic pathway [27]. Interestingly, when the cells were depleted of Survivin (i.e., upon treatment with YM155; *red bars*), a marked increase in the relative levels of F1,6BP and DHAP was observed, while only minimal amounts of the metabolite formed in the next step of the glycolytic pathway (i.e., glyceraldehyde 3-phosphate (G3P)), as well as those generated further downstream, were detected.

The accumulation of DHAP and loss of G3P in cells depleted of Survivin prompted us to focus on this step in the glycolytic pathway. The enzyme responsible for catalyzing the conversion of DHAP to G3P is TPI [31], and therefore we considered whether reductions in its expression could account for the differences in the intermediates detected under conditions when Survivin expression was blocked. However, the expression levels of TPI in cells treated with YM155 were nearly identical to the those detected in untreated cells (Fig. 3e). We then examined whether TPI might interact with Survivin. Immunoprecipitation experiments performed on lysates collected from cells ectopically expressing V5-tagged Survivin (Fig. 3f, *panels labeled whole cell lysates; WCL*) showed that TPI can be immunoprecipitated with Survivin (Fig. 3f, *panels labeled immunoprecipitation; IP*). An interaction between Survivin and TPI was similarly detected in MEFs ectopically expressing V5-tagged Survivin (*SI Appendix,* Fig. S2a).

Cells that expressed Survivin (i.e., control cells) exhibited high levels of TPI activity, whereas YM155 treatment resulted in a clear reduction in this reaction (Figure 3g, *compare bars 1 and 2*). The addition of purified recombinant Survivin to extracts collected from PDAC cells depleted of Survivin due to YM155 treatment then completely restored TPI activity to the levels observed in control cells (Figure 3g, *compare bars 2 and 3*). We further showed that adding recombinant Survivin to lysates collected from MEFs (which lack Survivin expression) gave rise to an ∼50% increase in TPI activity (*SI Appendix,* Fig. S2b).

Purified recombinant forms of Survivin and TPI were prepared (Figure 3h; *lanes labeled Input*) to determine whether these two proteins could directly interact and increase TPI activity. Fig. 3h shows that Survivin did not precipitate with a His-tagged form of TPI using nickel beads (*lanes labeled Precipitation*). Moreover, combining recombinant Survivin and TPI failed to stimulate TPI activity beyond the levels achieved when assaying the activity of recombinant TPI alone (Fig. 3i). Therefore, these findings indicated that the ability of Survivin to interact with TPI and promote its activation involves an additional protein(s).

### Microtubules are required for Survivin to promote TPI activity

One protein known to associate with both Survivin and TPI is tubulin [32–34]. α- and β-tubulin form heterodimers that further assemble into hollow, cylindrical tubes or microtubules, which typically provide cells with structural support and promote intracellular vesicle trafficking and mitosis [35]. Microscopy images obtained from immunofluorescence experiments performed on MIA PaCa-2 and PANC-1 cells using conditions that that specifically preserve cytoskeletal architecture, including microtubules (see *Materials and Methods* for details), show that both Survivin and TPI frequently localize together with tubulin within the cytosol in distinct ring-like structures (Figs. 4a and 4b and *SI Appendix,* Figs. S3a and S3b). Indeed, these structures were present in nearly 70% of the MIA PaCa-2 cells and PANC-1 cells (Fig. 4c). A closer examination showed that the rings consisted of all three proteins exhibiting a high degree of overlap (Figs. 4a and 4b, graphs under images, with a full co-localization analysis shown in *SI Appendix,* Figs. S4a and S4b) with diameters between 3-7 μm. The unique shape, size, and location of the structures consisting of Survivin, TPI and microtubules detected in PDAC cells are highly reminiscent of invadosome rosettes [36, 37], dynamic structures characterized by the presence of dense rings of filamentous actin (F-actin), together with various signaling proteins, adhesion molecules, and protein scaffolds that promote cell attachment and extracellular matrix degradation, resulting in increased cancer cell migration and invasion [36–38]. We show in Figure 4d that the ring-like structures detected in PDAC cells contained the canonical invadosome/podosome marker tyrosine substrate with five SH3 domains (TKS5) [38]. To further characterize these structures, we stained PDAC cells with rhodamine 565-conjugated phalloidin to detect F-actin, as well as with a tubulin antibody. The resulting images show that large rings of actin could be readily detected in the cells, providing additional evidence that these structures are invadosome rosettes (Figure 4e, row of images labelled *Control*). The rosettes of tubulin and actin were in close proximity but did not exactly overlap, with the tubulin rings encompassing the actin rings (Fig. 4e, images and corresponding graphs labelled *Control*, and *SI Appendix,* Fig. S5a). Consistent with previous findings [39, 40], we showed that the disruption of microtubules using the depolymerization agent nocodazole leads to loss of the actin-based rings (Fig. 4e, images and corresponding graphs labelled *Nocodazole*, and *SI Appendix,* Fig. S5a). Depletion of Survivin using YM155 often resulted in an even more striking effect on the actin rings (Fig. 4e, images and corresponding graphs labelled *YM155*, and *SI Appendix,* Fig. S5a). Nocodazole treatment is likely less effective at disrupting the formation of the ring superstructure because the association of Survivin with microtubules is known to potently block the actions of this drug to depolymerize microtubules [33,34].

**Figure 4.**
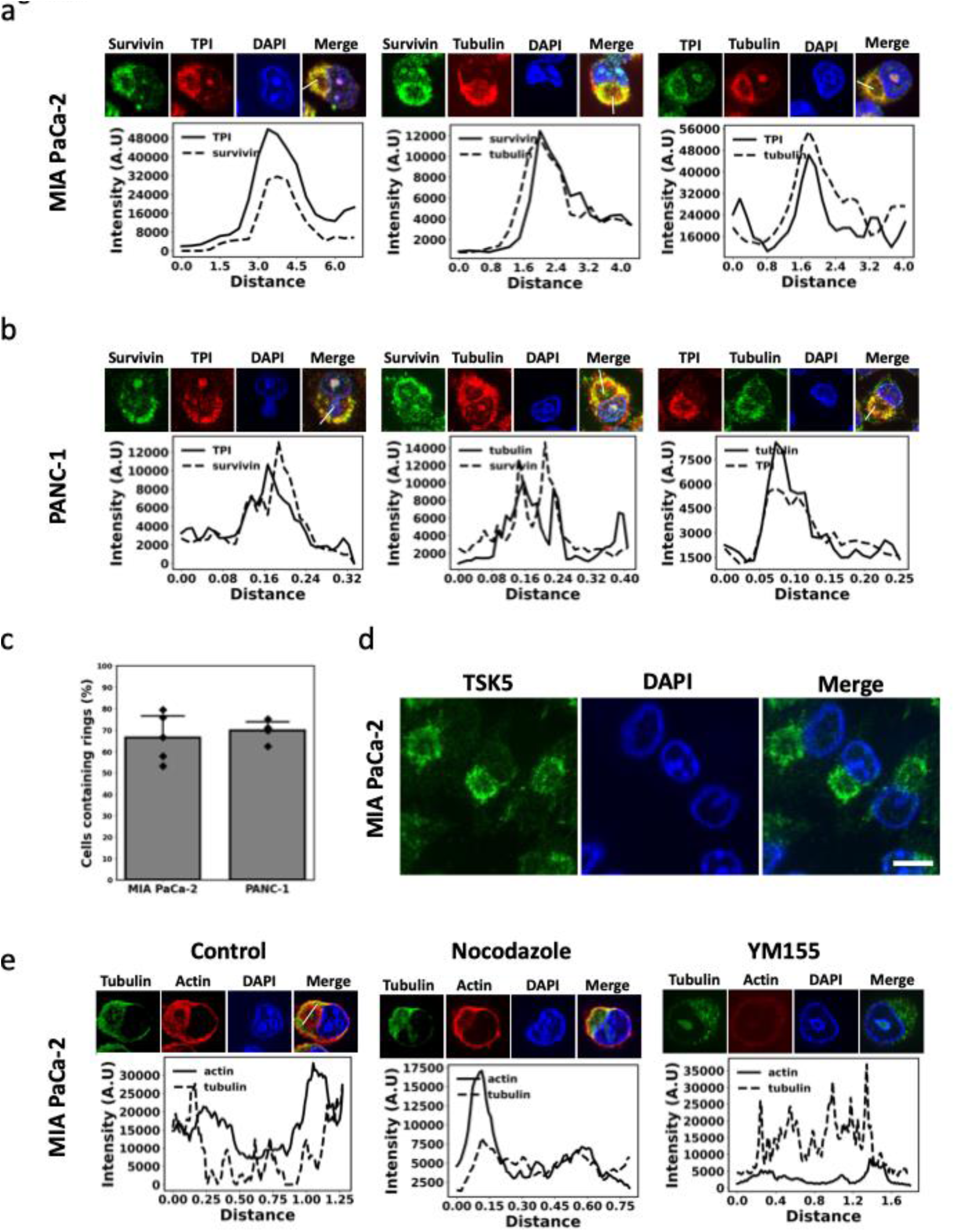
**a and b.** Fluorescent microscopy images of MIA PaCa-2 (a) and PANC-1 cells (b) immuno-stained for TPI, Survivin, and tubulin. Under each set of images the extent of co-localization of the proteins is shown, with the small lines in the merged images indicating where the analysis was performed. The full analysis of this experiment is shown in Supplementary Figs. 4a and 4b. **c.** The percentage of cells described in (a) and (b) with ring-like structures. **d.** Fluorescent microscopy images of MIA PaCa-2 cells immuno-stained for TSK5. **e.** Fluorescent microscopy images of MIA PaCa-2 cells treated without (*Control*; DMSO only) or with Nocodazole or YM155 and immuno-stained for tubulin and stained with rhodamine-conjugated phalloidin to visualize F-actin. Under each set of images the extent of co-localization of the proteins is shown, with the small lines in the merged images indicating where the analysis was performed. The full analysis of this experiment is shown in Supplementary Fig. 5a. All experiments shown were performed at least three independent times, and the cells in **a**, **b**, **d**, and **e** were also stained with DAPI to label nuclei.

Together these findings raised the possibility that microtubules were involved in the ability of Survivin to associate with TPI and promote its catalytic activity. To test this idea, MIA PaCa-2 cells were treated with the microtubule depolymerization agent nocodazole to determine the effects it has on the assembly of Survivin and TPI in the ring-like structures. While treating PDAC cells with nocodazole only partially disrupted the formation of the microtubule-based ring-like structures containing Survivin and TPI, YM155 treatment completely inhibited the assembly of these structures (Figs. 5a, row of images and corresponding graphs labelled *Nocodazole* and *YM155*, 5b, and *SI Appendix,* Fig. S5b). Nevertheless, nocodazole treatment resulted in a corresponding reduction in TPI activity (Fig. 5c). Collectively, these findings highlight that there is a tight correlation between increases in Survivin expression and the formation of invadosome rosette-like structures consisting of microtubules and TPI.

**Figure 5.**
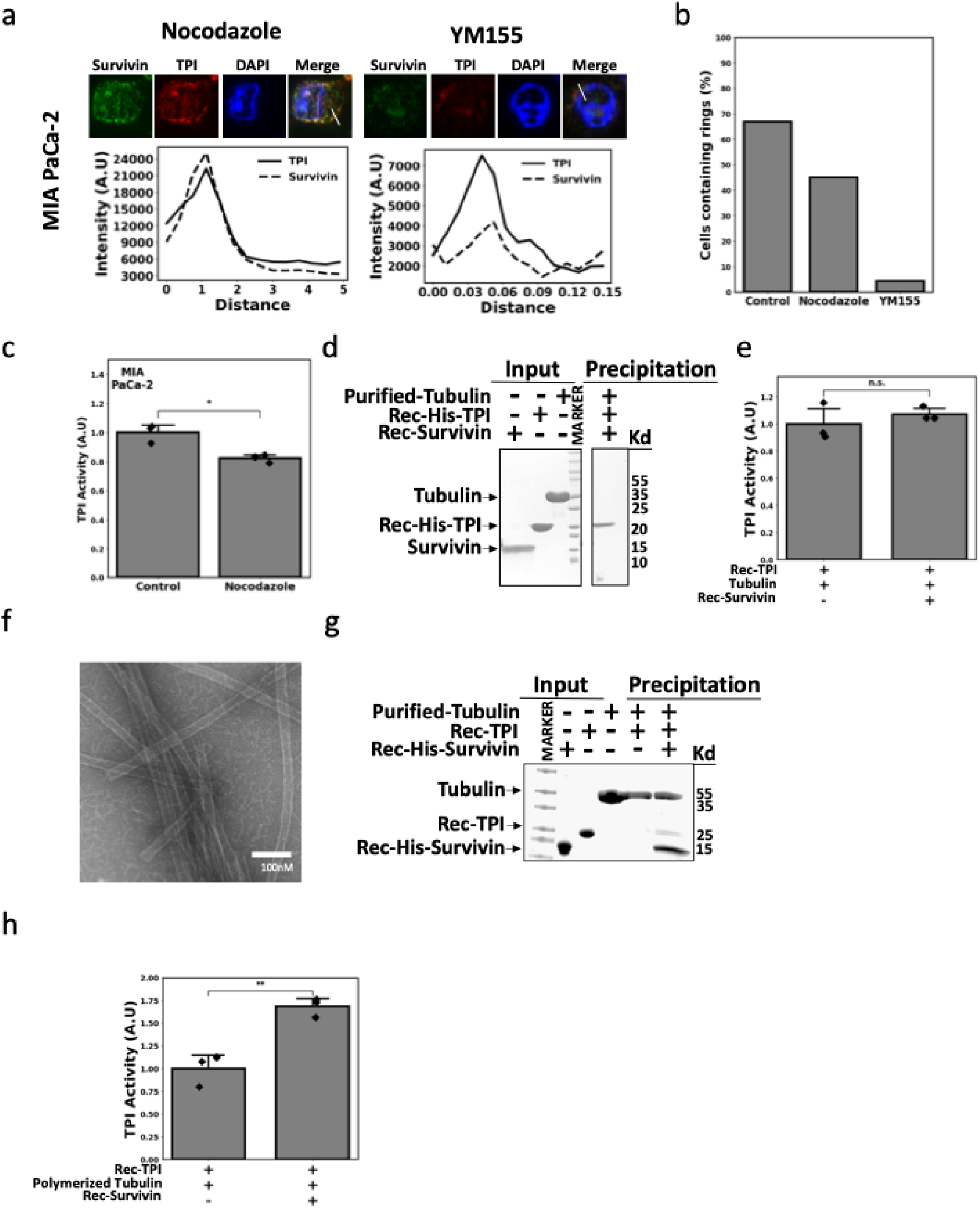
**a**. Fluorescent microscopy images of MIA PaCa-2 cells treated with nocodazole and YM155 and immuno-stained for TPI and Survivin. DAPI was used to label nuclei. Under each set of images the extent of co-localization of the proteins is shown, with the small lines in the merged images indicating where the analysis was performed. **b.** The percentage of cells described in (a) with ring-like structures. **c**. TPI activity assays were performed on MIA PaCa-2 cells treated without (*Control*; DMSO only) or with nocodazole. **d.** Protein precipitation experiments using nickel beads were performed on samples containing purified tubulin and recombinant forms of 6xHis-tagged TPI (Rec-His-TPI) and Survivin (Rec-Survivin). The proteins used in this experiment are shown (Input), as are the results of the precipitation assay (Precipitation) **e**. TPI activity assays were performed on the samples described in (d). **f.** Electron micrograph image showing tubulin incubated with GMPCPP polymerizes into microtubules. **g**. Protein precipitation experiments using nickel beads were performed on samples containing polymerized tubulin (i.e. microtubules) and recombinant forms of 6xHis-tagged TPI (Rec-His-TPI) and Survivin (Rec-Survivin). The proteins used in this experiment are shown (Input), as are the results of the precipitation assay (Precipitation). **h**. TPI activity assays were performed on samples described in (g). All experiments shown were performed at least three independent times. The data in **c, e** and **h** are presented as the mean, with p-values denoted as follows: n.s. not significant, * < 0.05, and ** < 0.01. Error bars indicate one standard deviation (SD).

Assays were then conducted to determine whether combining recombinant Survivin and TPI together with purified tubulin would result in the formation of a complex that could increase TPI activity. We initially performed these experiments using a non-polymerized form of tubulin, which exists predominantly as heterodimers of α- and β-tubulin (Fig. 5d, lanes labelled *Input*), and found neither tubulin nor Survivin precipitated with His-tagged TPI upon the addition of nickel beads (Fig. 5d, lanes labelled *Precipitation*), and that these conditions failed to boost TPI activity (Fig. 5e). The same experiments were then performed after the tubulin heterodimers were assembled into microtubules by the addition of the non-hydrolysable GTP analog, GMPCPP, which was confirmed by negative stain electron microscopy (Fig. 5f). When recombinant Survivin and TPI were combined with polymerized tubulin, i.e., microtubules, (Fig. 5g, lanes labelled *Input*), all three proteins were precipitated by His-tagged Survivin (Fig. 5g, lanes labelled *Precipitation*). TPI activity assays performed on these samples showed that a marked increase occurred in TPI, compared to samples that contained only TPI and microtubules (Fig. 5h).

### Survivin promotes the localization of additional glycolytic enzymes to microtubules

Our findings demonstrate that the high levels of Survivin expression associated with PDAC cells work together with microtubules to form a structure that can associate with TPI and increase it enzymatic activity. However, they also raise the question as to whether additional glycolytic enzymes might also be recruited to these unique Survivin-microtubule-base structures to generate a “hub” or metabolon that would increase the efficiency of the glycolytic pathway. Therefore, immunofluorescence was performed on MIA PaCa-2 and PANC-1 cells to determine whether the next enzyme in the glycolytic pathway after TPI, GAPDH [8,27], is similarly localized to the Survivin-microtubule-TPI complex. This is a particularly attractive idea to consider, since the reaction carried-out by GAPDH is a rate-limiting step in glycolysis [41]. The resulting microscopy images taken of the cells show that GAPDH does indeed associate with the same ring-like structures as TPI (Fig. 6a, images and corresponding graphs and *SI Appendix,* Fig. S6a, with a full co-localization analysis shown in *SI Appendix,* Fig. S6b), suggesting that Survivin, upon binding to microtubules, stimulates glycolytic activity in KRAS-driven pancreatic cancer cells by increasing the localized concentrations of glycolytic enzymes, including TPI and GAPDH. This ensures that the production of G3P from DHAP by TPI is rapidly converted by GAPDH to 1,3BPG, and as a result increases the level of glycolytic activity required for the rapid growth of cancer cells (Fig. 6b).

**Figure 6.**
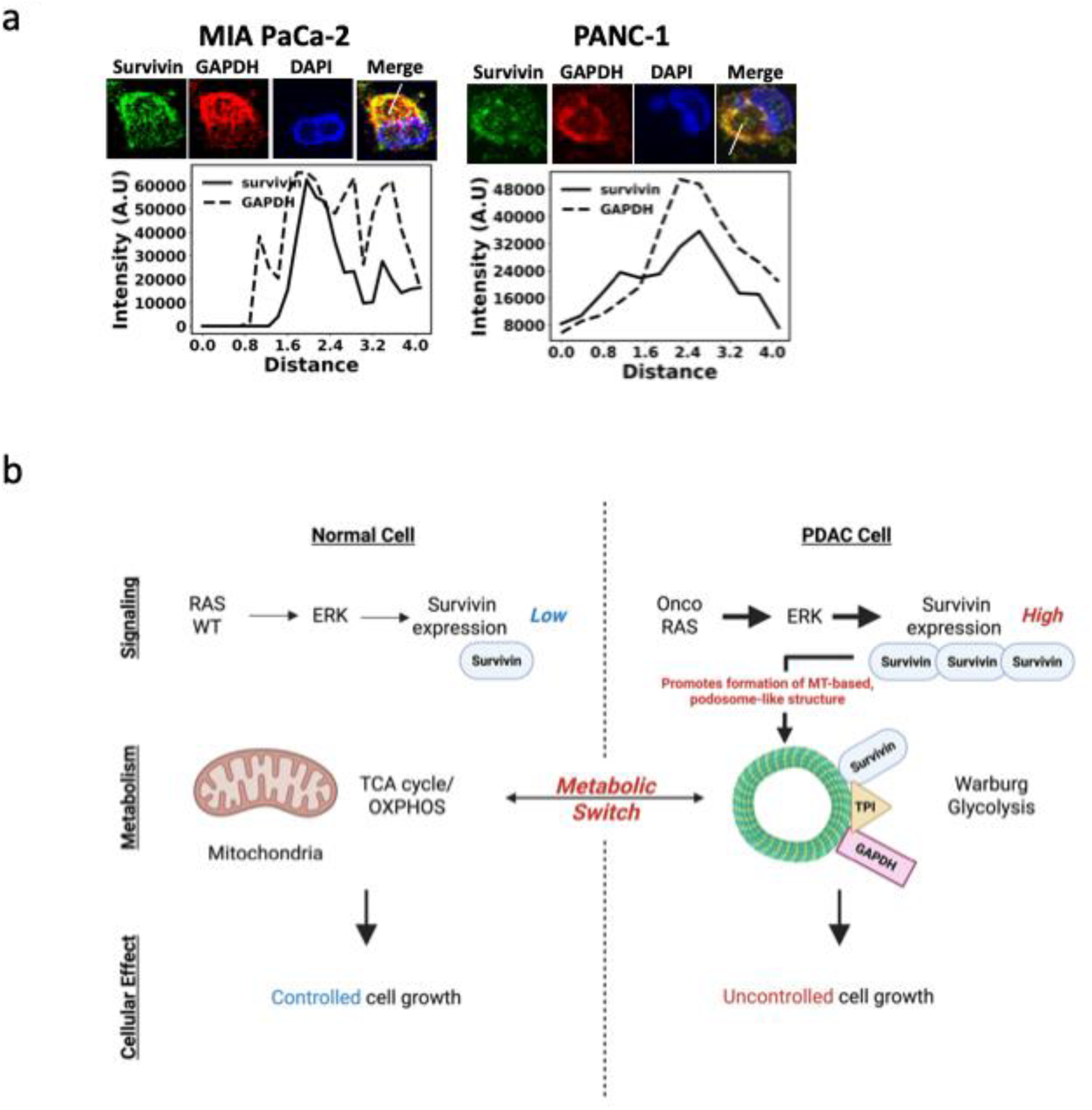
**a**. Fluorescent microscopy images of MIA PaCa-2 cells immuno-stained for Survivin and GAPDH. DAPI was used to label nuclei. Under each set of images the extent of co-localization of the proteins is shown, with the small lines in the merged images indicating where the analysis was performed. The full analysis of this experiment is shown in Supplementary Fig. 6b. The experiment shown was performed at least three independent times. **b**. Schematic showing how normal cells express Survivin at very low levels and typically undergo oxidative phosphorylation (OXPHOS), which takes place in mitochondria (left side). However, the expression of oncogenic forms of Ras in PDAC cells activates signaling proteins, including ERK, that results in the increased expression of Survivin. The high levels of Survivin associated with these cells promotes the assembly of microtubules into unique ring-like structures that further recruit glycolytic enzymes, including TPI and GAPDH, to promote Warburg glycolysis (right side).

## Discussion

Survivin has been implicated in promoting several cancer cell phenotypes, with its best-established roles being the inhibition of apoptosis and promotion of cell growth by inhibiting negative regulators of the cell cycle, increasing the expression of Myc, and ensuring the proper segregation of chromosomes into daughter cells during mitosis [15–24]. There have also been suggestions that Survivin contributes to the metabolic reprogramming of cancer cells [42]; however, how it plays such a role has remained unclear. We now identify a previously unappreciated connection between oncogenic forms of KRAS, Survivin, the cytoskeleton, and increases in glycolytic activity in PDAC cells and patient-derived organoids. We show that oncogenic KRAS mutants signal a robust increase in Survivin expression, which results in the formation of a unique complex between Survivin and microtubules. One of the key features of this complex is its large ring-like structure that bears a striking resemblance to invadosome rosettes [36,37]. These higher order structures composed of interconnected invadosomes are known to function as critical points of contact between the cell and its extracellular environment that promote cell attachment, migration and invasion. Invadosomes are characterized by dense actin-rich rings and here we now demonstrate that these actin-based structures can be encompassed by additional rings of Survivin and microtubules. While Survivin has been shown to interact with microtubules at stages of the cell cycle other than mitosis [33], its role in these contexts has remained largely unexplored. We now show the Survivin-microtubule complex that assembles in cells recruits multiple glycolytic enzymes, including TPI and GAPDH, that are essential for maintaining high levels of glycolytic activity in PDAC cells necessary for their ability to rapidly proliferate. These new findings significantly expand our understanding of Survivin and highlight its role as a key effector of oncogenic KRAS signaling, linking cytoskeletal re-organization to metabolic reprogramming. They also help to explain why proteomic analyses have identified several glycolytic enzymes on isolated invadosomes from cancer cells, with the idea being that they provide the energy needed to maintain their structure and function [43].

Given that Survivin expression is very low or undetectable in most differentiated adult cells and tissues [15, 16], its interaction with microtubules that leads to the assembly of a glycolytic enzyme complex essential for tumor progression offers new therapeutic possibilities by targeting this metabolic hub which can bypass the challenges in directly targeting Survivin [15]. Both microtubules and glycolysis have been targeted for decades using drugs like paclitaxel and 2-deoxyglucose (2-DG) [44, 45] and shown some promise for cancer treatment. However, our findings suggest that disrupting their point of intersection within the Survivin–TPI–microtubule complex described here may offer a more precise and effective therapeutic strategy and thus represent a metabolic vulnerability, or “Achilles’ heel,” in KRAS-driven cancers.

## Materials and Methods

### Cell Culture

All cell lines were obtained from the ATCC and maintained at 37°C in a humidified incubator with 5% CO₂. The BxPC-3, PANC-1, MIA PaCa-2, HEK293T, MDA-MB-231, and mouse embryonic fibroblasts (MEFs) were cultured in growth medium: Dulbecco’s Modified Eagle’s Medium (DMEM; Thermo-Fisher Scientific, Cat #11965092) supplemented with 10% fetal bovine serum (FBS; Thermo-Fisher Scientific, Cat #16010159), while NIH3T3 mouse embryonic fibroblasts (MEFs) were grown in DMEM containing 10% calf serum (CS; Thermo-Fisher Scientific, Cat #A5256701).

### PDAC Organoid Culture

Patient-derived organoids were obtained from Dr. David Tuveson (Cold Spring Harbor Laboratory). hM1A, hT105, PT6, and PT8 human PDAC patient-derived organoids were cultured in growth factor reduced Matrigel (Corning, Cat #CB-40234A) domes in complete human feeding medium: Advanced DMEM/F12 (Thermo-Fisher Scientific, Cat #11320033) based L-WRN conditioned medium (ATCC, Cat #CRL-3276) supplemented with B27 supplement (Thermo-Fisher Scientific, Cat #17504044), 10 mM HEPES (Thermo-Fisher Scientific, Cat #11344041), 0.01 μM GlutaMAX (Thermo-Fisher Scientific, Cat #35050061), 10 mM nicotinamide (Sigma-Aldrich, Cat #N0636), 50 ng/mL hEGF (Thermo-Fisher Scientific, Cat #AF-100-15), 100 ng/mL hFGF10 (Thermo-Fisher Scientific, Cat #AF-100-26), 0.01 μM hGastrin I (TOCRIS, Cat #3006), 500 μM A83-01 (TOCRIS, Cat #2939), and 1.25 mM (hM1A, hT105), or 1 mM (PT6 and PT8) N-acetylcysteine (Sigma-Aldrich, A9165). The media for PT6 and PT8 additionally contained 10 μM SB202190 (Sigma Aldrich, Cat #S7076). 10.5 μM Y27632 (Selleckchem, Cat #S1049) was added for the first two days after reseeding the cells. The known genetic mutations associated with each of the organoids are listed in Fig. 2a.

### Cell Transfection

Cells were grown to 50–70% confluency before being transfected with the indicated expression constructs using FuGene HD (Promega, Cat #E2311), according to the manufacturer’s instructions. Four hours after the transfection, the medium was replaced with growth medium, and the cells were allowed to grow for an additional 36 hours before they were collected.

### Virus Production and Infection of Cells

Lentivirus was produced by transfecting HEK293T cells with either BIRC5-targeting shRNA or a scrambled shRNA control, together with the viral packaging constructs pCMV delta R8.2 (Addgene, Cat #12263) and pMD2.G (Addgene, Cat #12259). The virus-containing medium was collected 24- and 48-hours later and combined with polybrene (10 μg/mL, Sigma Aldrich, Cat #H9268) before being used for cell transduction.

### Cell and Organoid Inhibitor Treatments

Cultures of cells and organoids maintained in their growth medium were treated for the indicated lengths of time with DMSO, 100 nM YM155 (unless indicated otherwise; MedChemExpress, Cat #HY-10194), 5.0 mM 2-DG (unless indicated otherwise; MedChemExpress, Cat #HY-13966), 1.0 μM AMG510 (MedChemExpress, Cat #HY-114277), 1.0 μM SCH772984 (MedChemExpress, Cat #HY-50846), and 1.0 μM nocodazole (Sigma Aldrich, Cat #M1404).

### Protein Expression and Purification

pET-28a (+) bacterial expression plasmids encoding His-tagged forms of TPI or Survivin were generated (Twist Bioscience), transformed into *E. coli*, and grown at 37°C in LB broth until the cultures reached an optical density (OD) of 0.6 at 600 nm (A600). The temperature was then reduced to 18°C, and protein expression was induced by the addition of 100 μM IPTG for 16 hours. The bacterial cultures were pelleted by centrifugation, resuspended in bacterial lysis buffer (50 mM Tris, pH 8.0, 500 mM NaCl), and the mixture was sonicated before being centrifuged at 185,000 × g for 45 minutes. The supernatant was incubated with nickel beads for 20 minutes at 4°C, after which the beads were loaded onto a gravity-flow column. The beads were rinsed using wash buffer (50 mM Tris, pH 8.0, 500 mM NaCl, and 20 mM imidazole), and the protein was eluted from the beads with wash buffer containing 250 mM imidazole. The His-tag was removed from the recombinant proteins using thrombin, and then the samples were loaded onto a 5 mL HiTrap His column connected to a 5 mL Q HP column (both from Cytiva Life Sciences). Proteins were eluted using a gradient between Buffer A (bacterial lysis buffer) and Buffer B (bacterial lysis buffer with 1 M NaCl). The buffer of the purified protein was exchange into BRB80 buffer (100 mM PIPES, 1 mM MgCl₂, 1 mM EGTA, pH 7.2) using dialysis tubing with a 3.5 kDa molecular weight cutoff (ThermoFisher Scientific, cat #88242). The recombinant form of Survivin was concentrated to 3 mg/mL, and recombinant TPI was concentrated to 5 mg/mL, before being aliquoted, snap-frozen, and stored at −80°C. For some experiments the His-tag on the recombinant proteins was not removed.

### Western blot Analysis

Cells and organoids grown and treated as indicated, were lysed using cell lysis buffer (25 mM Tris, 100 mM NaCl, 1.0 mM EDTA, 1 mM DTT, 1 mM β-glycerol phosphate, 1% Triton X-100, 1 μg/mL aprotinin, and 1 μg/mL leupeptin) and protein concentrations were determined using the Bradford assay (Bio-Rad, Cat #5000006). Equal concentrations of protein lysates were combined with Laemmli sample buffer and resolved on 4–20% gradient SDS-PAGE gels (ThermoFisher Scientific, Cat #XP04200BOX), and transferred to 0.45 μm PVDF membranes (ThermoFisher Scientific, Cat #88518). The membranes were blocked with 10% bovine serum albumin (Sigma Aldrich, Cat #A9647) in TBST buffer (20 mM Tris, 2.7 mM KCl, 140 mM NaCl, and 0.5% Tween-20) and incubated overnight at 4°C with one of the following primary antibodies diluted 1:1000 in TBST: ERK (Cell Signaling Technology, Cat #4695), phospho-ERK (Cell Signaling Technology, Cat #9101), Survivin (Cell Signaling Technology, Cat #2808), TPI (Cell Signaling Technology, Cat #34088), tubulin (Cell Signaling Technology, Cat #2148), V5-tag (Cell Signaling Technology, Cat #13202), Myc-tag (Cell Signaling Technology, Cat #2276), GAPDH (Cell Signaling Technology, Cat #2118) and Vinculin (Cell Signaling Technology, Cat #13901). Following primary antibody incubation, membranes were washed and incubated for 1 hour at room temperature with a horseradish peroxidase (HRP)-conjugated secondary antibody (Cell Signaling Technology, Cat #7074). The blots were exposed to ECL reagent and developed using either HyBlot CL Autoradiography Film (Thomas Scientific, Cat #1141J52) with a Konica Minolta SRX-101A developer or imaged using a ChemiDoc MP system (Bio-Rad).

### Cell Growth Assay

Cells plated in 6-well dishes at a density of 30,000 cells per well were treated as indicated. After 48 hours, the cells were detached using trypsin and counted using a TC-20 automated cell counter (Bio-Rad). The growth of the untreated cells was normalized to 1-fold, and the fold change in growth of the treated cells was compared to this baseline.

### Lactic Acid Assay

Cells plated in 12-well dishes at a density of 50,000 cells per well were treated as indicated for 10 hours, at which point, the conditioned medium was collected and filtered using a 3 kDa molecular weight cut off spin column (Amicon, Cat #UFC800304). The lactic acid levels in the medium were determined using the Colorimetric L-Lactate Assay Kit (Abcam, Cat #ab65330) according to the instructions provided by the manufacturer. Absorbance was measured at 450 nm using the TECAN Spark microplate reader (TECAN).

For measuring lactic acid production via NMR, 10⁵ cells were seeded in 12-well plates and left settled overnight. The cells were then washed with PBS, and 750 μL of fresh experimental medium (low-glucose DMEM, Thermo Fisher Cat #11885-084, or RPMI, each containing 10% FBS) was added for 24 hours. The conditioned media were then harvested and filtered using Amicon Ultra 10 kDa MWCO units. A 400 μL filtered conditioned medium was mixed with 200 μL of 99% D₂O containing 6 mM TSP and 3 mM imidazole. The NMR spectra were recorded using a Bruker Advance III 700 MHz Spectrometer, and lactic acid was profiled and quantified by Chenomx.

### TPI Activity Assay

TPI activity was determined using the colorimetric TPI Activity Assay Kit (Abcam, Cat #ab197001). Briefly, cells treated as indicated for 10 hours were detached from the plate using trypsin and counted using a TC-20 automated cell counter (Bio-Rad). One million cells were pelleted and lysed using the buffer provided in the kit. The lysates were then centrifuged at 12000 rpm for 5min, and the supernatants were transferred to a 96-well plate, and their fluorescence was measured at 450 nm using the TECAN Spark microplate reader. In some cases, different combinations of recombinant proteins and purified tubulin (Cytoskeleton, Cat #T240) were assayed for TPI activity using the same approach.

### Immunofluorescence

Cells plated on glass coverslips were treated as indicated for 12 hours then briefly extracted in BRB80 containing 4mM EGTA and 0.5% Triton x-100 followed by the either the addition of a 1:1 mixture of ice cold methanol (ThermoFisher Scientific, Cat #268280010) and acetone (Thermo-Fisher Scientific, Cat #AA22928K2) for 5 minutes at −20C, washed with Tris buffered saline (TBS) with 0.1% Triton X-100 (TBST) or the addition of BRB80 with 4mM EGTA and 0.5% glutaraldehyde, then quenched in PBS with 0.1% sodium borohydride. The coverslips were blocked using TBST with 5% BSA for 1 hour, followed by incubation at room temp for 1 hour with primary antibodies against Survivin (Invitrogen, Cat #MA5-17035 or Cell Signaling Technology, Cat #2808), tubulin (Cell Signaling Technology, Cat #2148), V5-tag (Cell Signaling Technology, Cat #13202), Myc-tag (Cell Signaling Technology, Cat #2276), tubulin (ThermoFisher, Cat #MA1-19400), GAPDH (Cell Signaling Technology, Cat #2118), TPI (Cell Signaling Technology, Cat #34088), or TKS5 (ThermoFisher, Cat# PA5-117156). The coverslips were again washed in TBST before being incubated in fluorescent-conjugated secondary antibodies, rhodamine-conjugated phalloidin (Thermo-Fisher Scientific, Cat #R415), and/or DAPI (Thermo-Fisher Scientific, Cat #62248). Imaging was performed on a Keyence BZ-X810 inverted fluorescence phase-contrast microscope (Keyence Corp, Cat #BZPA60). The colocalization tests were performed using the built in colocalization tool with auto thresholding in Image J.

### Transmission Electron Microscopy

Formvar/carbon copper grids were plasma cleaned before applying 4 μL of the Survivin/TPI/Tubulin complex. After a 30-second incubation, the excess liquid was wicked away using Whatman paper. The grids were then stained with 2% uranyl acetate for 30 seconds, and the staining step was repeated. Following staining, the grids were allowed to air dry for 5 minutes before imaging on a Talos F200i transmission electron microscope (TEM) at 140 kV.

### Microtubule Formation

Porcine tubulin (Cytoskeleton Inc., Cat #T240) was resuspended in BRB80 buffer containing 1 mM GTP, aliquoted, and flash-frozen at a concentration of 10 mg/mL. Microtubule seeds were generated using an aliquot of tubulin diluted in BRB80 to 2.5 mg/mL followed by the addition of 1mM GMPCPP and 1mM DTT. The mix was then warmed to 37C for 5min then spun at 350,000 × g for 5min, the supernatant was cleared, and the pellet was washed with warm buffer before being resuspended in BRB80. For experiments, an aliquot was thawed and diluted to 2.5 mg/mL in BRB80 buffer, 0.75 mg/mL recombinant Survivin, and 1.0 mg/mL recombinant TPI were combined with microtubule seeds at a 1:20 ratio. The solution was incubated at 37°C for 15 minutes to allow for microtubule polymerization. The mixture was centrifuged at 350,000 × g for 5 minutes to pellet the microtubules that formed, and the resulting pellet was washed twice with BRB80 buffer at 37°C and then was resuspended in room temp BRB80. Samples were visualized by electron microscopy as described above.

### Immunoprecipitations

Lysates of cells ectopically expressing V5-tagged Survivin or the vector alone (1600 μg of each lysate) were incubated with anti-V5 affinity beads (Sigma Aldrich, Cat #SAE0203) at 4°C for 3 hours on a rotor. The beads were washed three times with cell lysis buffer, resuspend in Laemmli sample buffer and boiled for 5 minutes. The samples were subjected to Western blot analysis.

### Protein Precipitation Assay

Recombinant His-tagged TPI was combined with different combinations of recombinant Survivin and purified tubulin to final concentrations of 0.05 mg/mL, 0.15 mg/mL, and 0.25mg/ml respectively. The samples were incubated at RT for 30 minutes, before nickel beads (Gold Bio-Technology, Cat #H-351-2) were added to the mixture and incubated for an additional 30 minutes. The beads were washed three times with BRB80 plus 0.025% triton X 100, resuspended in Laemmli sample buffer and boiled for 5 minutes. The samples were resolved on 4–20% gradient SDS-PAGE gels, and stained with Aquastain protein gel plus (Bulldog Bio).

### Glucose Uptake Assay

Cells plated on glass-bottom microwell dishes were treated as indicated for 10 hours, at which point the media was replaced with fresh growth media containing 100 μM of the fluorescent glucose analog 2-NBDG (ThermoFisher Scientific, Cat #N13195). Thirty minutes later the cells were washed with PBS and fixed with 4% paraformaldehyde. The microwell dishes were Imaged using the Keyence microscope at 63× magnification, and fluorescence intensity was quantified using ImageJ software.

### LC-MS/MS

Cells treated without (DMSO) or with 500nM for 10 hours YM155 were snap-frozen in liquid nitrogen, and metabolites were extracted using HPLC-grade methanol (Thermo-Fisher Scientific, Cat #325740025). Pyruvate was derivatized using an established protocol. Targeted metabolite analysis was performed using a Vanquish Horizon UHPLC system (Thermo-Fisher Scientific) coupled to a TSQ Quantis triple quadrupole mass spectrometer equipped with a HESI ion source (Thermo-Fisher Scientific). Standard compounds, including D-fructose-6-phosphate (F6P; Sigma Aldrich, Cat #F3627), D-fructose-1,6-bisphosphate (F1,6BP; Sigma Aldrich, Cat #F6803), dihydroxyacetone phosphate (DHAP; Sigma Aldrich, Cat #D7137), DL-glyceraldehyde-3-phosphate (G3P; Sigma Aldrich, Cat #17865), D-(-)-3-phosphoglyceric acid (3PG; Sigma Aldrich, Cat #P8877), and sodium pyruvate (Thermo-Fisher Scientific, Cat #890-1840IL) were used as reference standards. For chromatographic separation of non-pyruvate metabolites, the mobile phase consisted of 20% HPLC-grade water (Thermo-Fisher Scientific, Cat #022934), 80% HPLC-grade acetonitrile (Thermo-Fisher Scientific, Cat #022927.K2), and 0.1% (v/v) formic acid (Thermo-Fisher Scientific, Cat #A117-50) as Mobile Phase A, while Mobile Phase B contained 70% acetonitrile, 30% water, and 0.1% formic acid. Metabolites were separated using a XBridge Amide column (100 × 2.1 mm i.d., 3.5 μm; Waters). The mass spectrometer was operated in negative ion mode with a spray voltage of 2.5 kV, an ion transfer tube temperature of 350°C, and a vaporizer temperature of 400°C. The sheath gas, auxiliary gas, and spare gas were set to 60, 15, and 2 arbitrary units, respectively. The mass-to-charge ratio (m/z) was analyzed within a range of 100–400 or 80–200, and metabolite identification was validated using synthetic standards. For pyruvate-specific analysis, chromatographic separation was performed using an Agilent InfinityLab Poroshell 120 EC-C8 column (50 × 2.1 mm i.d., 2.7 μm; Agilent, Cat # #699775-902) at 50°C. The mobile phase consisted of 99.9% water with 0.1% formic acid (Mobile Phase A) and 99.9% acetonitrile with 0.1% formic acid (Mobile Phase B). The mass spectrometer was operated in negative ion mode with a spray voltage of 2.5 kV, an ion transfer tube temperature of 350°C, and a vaporizer temperature of 325°C. Sheath gas, auxiliary gas, and spare gas were set to 50, 10, and 1 arbitrary units, respectively, with an m/z range of 200–400. Derivatized pyruvate was included as a positive control. All LC-MS/MS data were analyzed using the XCalibur software suite (Thermo-Fisher Scientific).

## Quantification and Statistical Analysis

All experiments were performed at least three independent times. Statistical significance of the experiments was determined using Student’s t-tests, and the data was presented as means +/-standard deviations (SD). The relative mRNA levels in PDAC and normal tissue samples were determined and graphed using the TMNPlot online software.

## Author Contributions

JN, W-HC, RAC, and MAA conceived the idea, designed, and performed experiments, and wrote the manuscript. SEA, MRZ, TM, RY, JRL, SE, HHL, MTL, ELJ, and KLB performed experiments.

## Author Approvals

All Authors have seen and approved this manuscript. This work has not been published or accepted for publication in any journals at this time.

## Classification

Biological Sciences, Cell Biology

