## Supplemental Figures for "Survivin Promotes the Formation of a Microtubule-Based Glycolytic Hub"

**Figure S1**

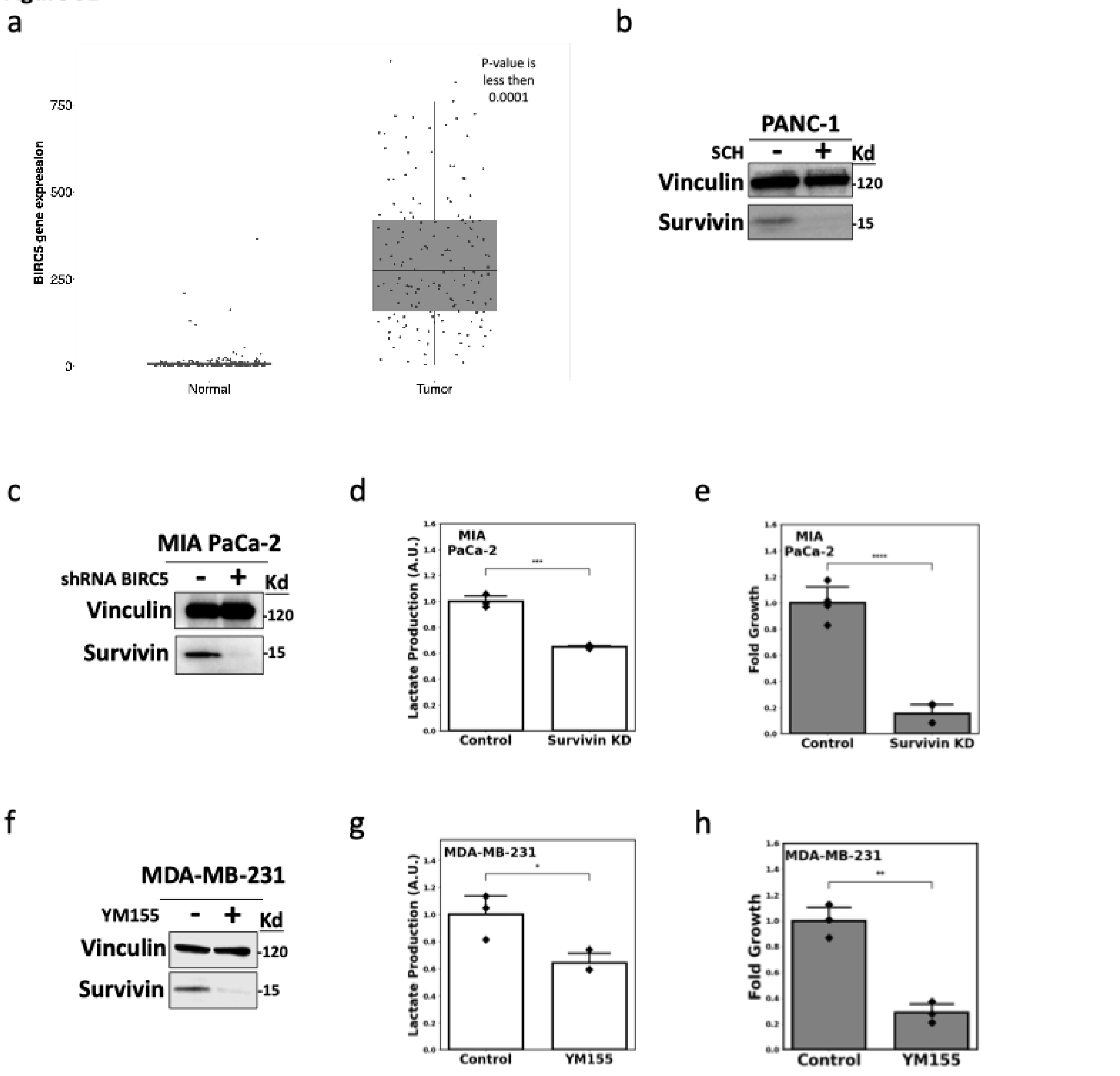


**Figure S1.** a. Relative BIRC5 transcript levels (which encodes the Survivin protein) in normal pancreatic tissue (*Normal*) and PDAC samples (Tumor). Data was generated using GEPIA 2 and the TCGA database, and the p value is indicated in the graph. b. Western blot showing Survivin expression levels in PANC-1 cells treated without (-; DMSO only) or with (+) SCH772984 (SCH). c. Western blot showing Survivin expression levels in MIA PaCa-2 cells expressing control shRNA (-) or BIRC5 targeting shRNA (+). d and e. Lactic acid (d) and growth assays (e) were performed on the cells treated as described in (c).

f. Western blot showing Survivin expression levels in triple-negative MDA-MB-231 breast cancer cells treated without (-; DMSO only) or with (+) 100 nM YM155. g and h. Lactic acid (g) and growth assays (h) were performed on MDA-MB-231 cells treated as described in (f). All experiments shown were performed at least three independent times, and vinculin was used as a loading control for the Western blots. The data in d, e, g, and h are presented as the mean, with p-values denoted as follows: * < 0.05, ** < 0.01, *** < 0.001, **** < 0.0001. Error bars indicate one standard deviation (SD).

**Figure S2**

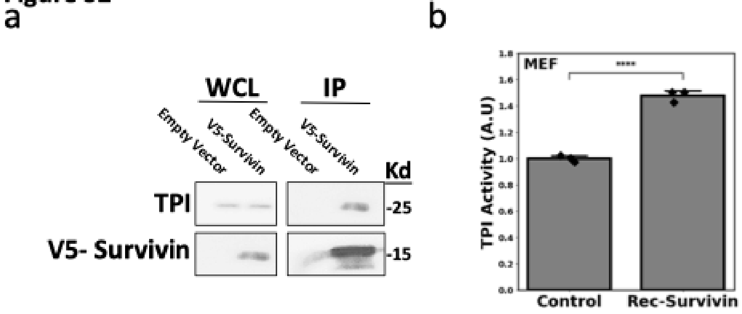


**Figure S2.** a. Western blot showing TPI and V5-tagged Survivin expression levels in the whole-cell lysates (WCL), and immunoprecipitations performed on these lysates using a V5 antibody (IP), from MEFs transfected with either an empty vector or a vector encoding V5-tagged Survivin. b. TPI activity assays were performed on the lysates collected from MEFs (*Control*), or after the sample lysates were incubated with a recombinant form of Survivin (Rec-Survivin). All experiments shown were performed at least three independent times, and TPI was used as a loading control for the Western blots. The data in b are presented as the mean, with p-values denoted as follows: **** < 0.0001. Error bars indicate one standard deviation (SD).

Figure S3

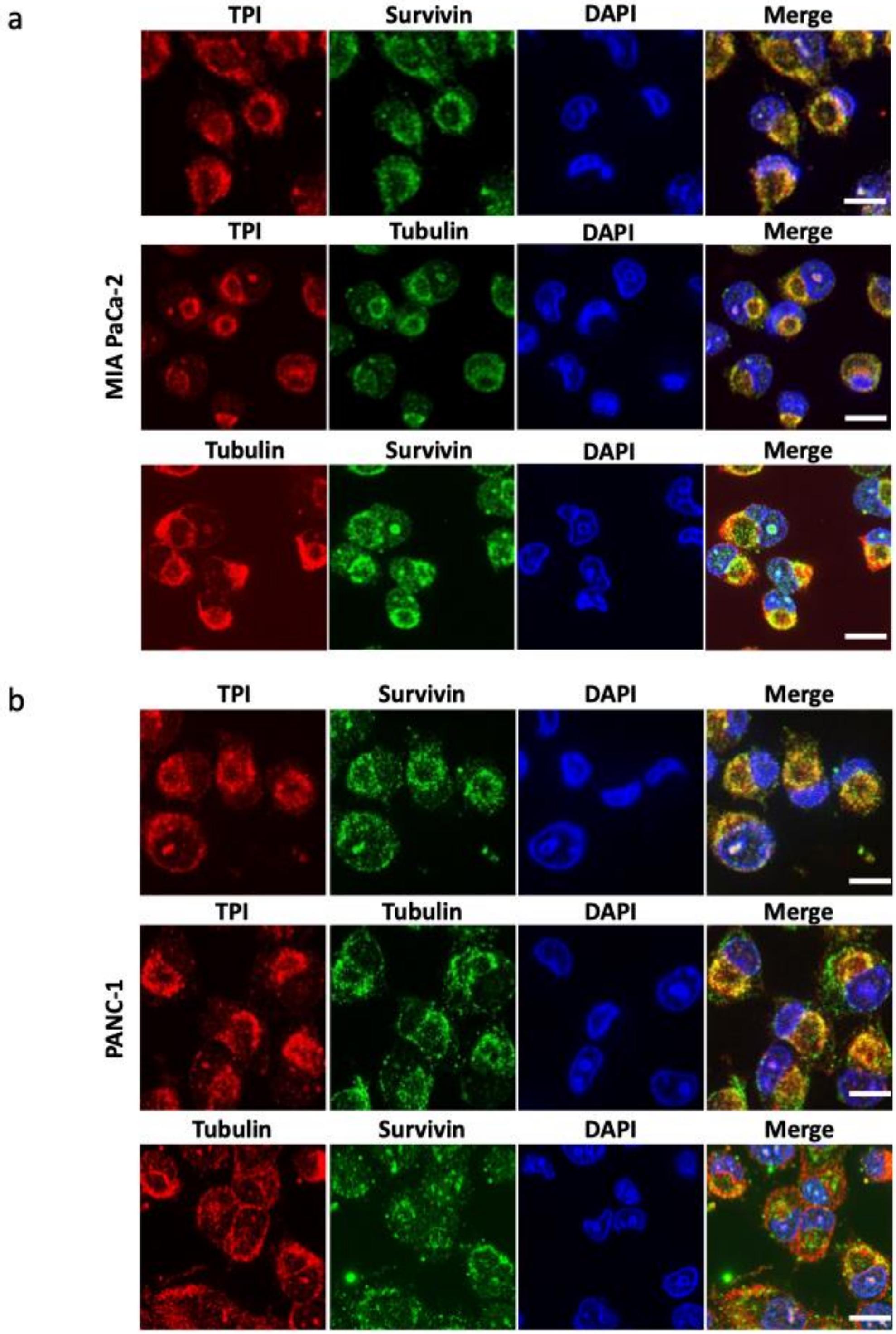


**Figure S3.** a and b. Full-sized fluorescent microscopy images of MIA PaCa-2 (a) and PANC-1 cells (b) immuno-stained for TPI, Survivin, and tubulin. The cells were also stained with DAPI to label the nuclei. These images were used to generate the images of cells shown in Figs. 4a and 4b. All experiments shown were performed at least three independent times.

**Figure S4**

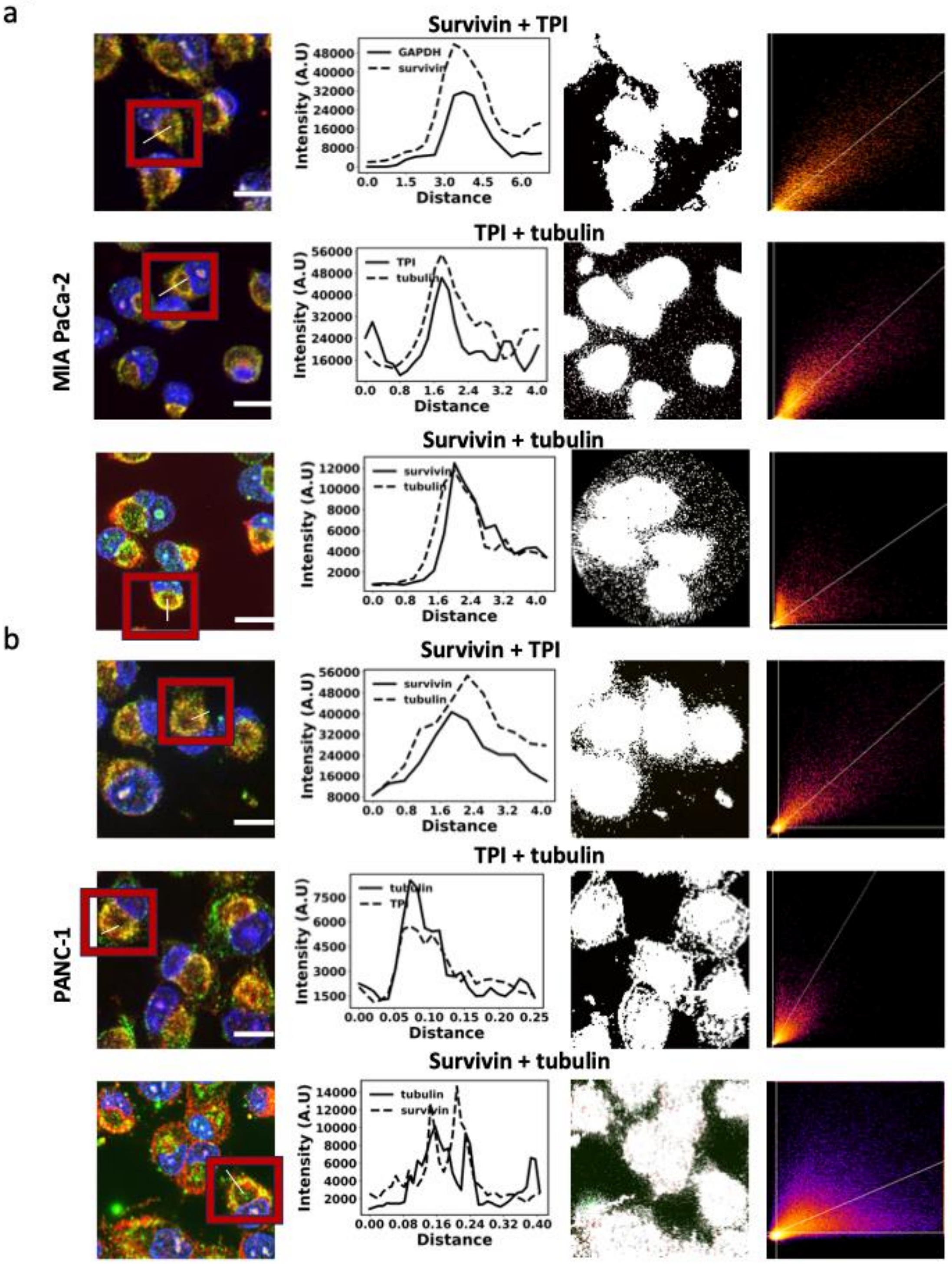


**Figure S4.** a and b. The full co-localization analyses performed on the merged images of the cells shown in Figs. 4a and 4b using ImageJ. The boxed cell shown in each of the first panels was analyzed for the co-localization of the two indicated proteins, with the white line indicating where the analysis was performed. The second panel in each set are graphs showing the extent of co-localization of the two proteins, while the third and fourth panels show the co-localization mask and the co-localization analysis, respectively. c. Fluorescent microscopy images of cells immunostained for TKS5. The cells were also stained with DAPI to label the nuclei. All experiments shown were performed at least three independent times.

Figure S5

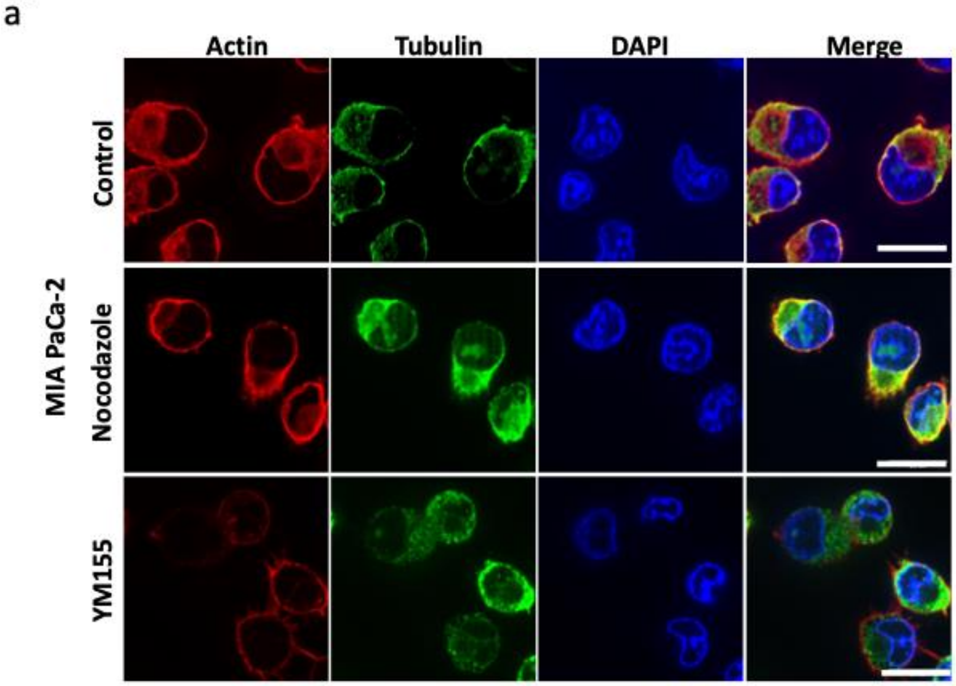


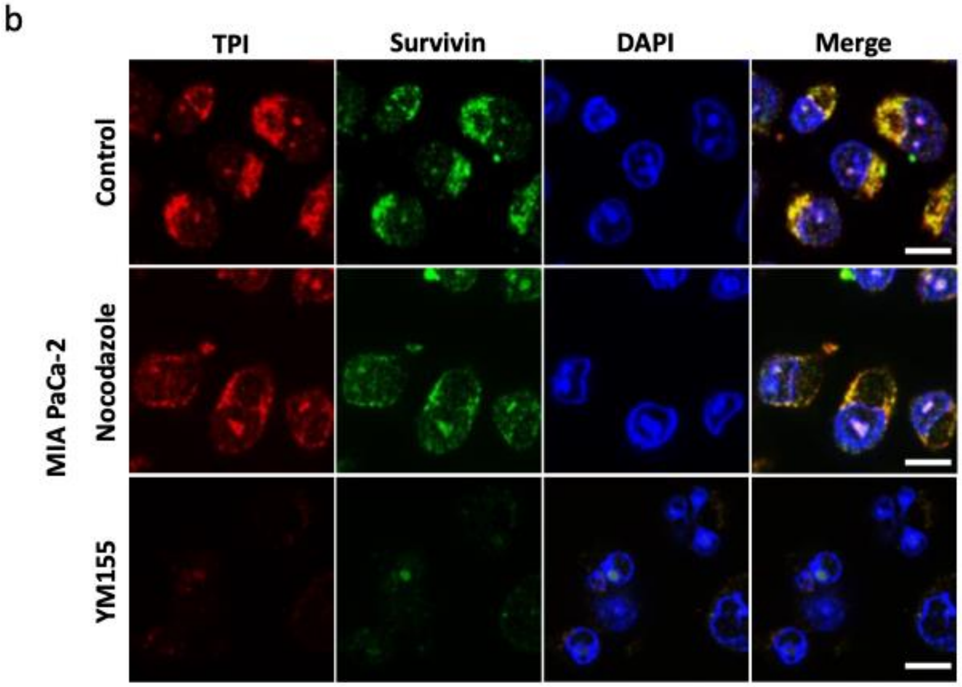


**Figure S5 a.** Full-sized fluorescent microscopy images of MIA PaCa-2 cells treated without (*Control*; DMSO only) or with Nocodazole and YM155 and immuno-stained for F-actin and tubulin. The cells were also stained with DAPI to label the nuclei. These images were used to generate the images of cells shown in Fig. 4d. b. Full-sized fluorescent microscopy images of MIA PaCa-2 cells treated without (*Control*; DMSO only) or with Nocodazole and YM155 and immuno-stained for TPI and Survivin. The cells were also stained with DAPI to label the nuclei. These images were used to generate the images of cells shown in Fig. 5a.

**Figure S6**

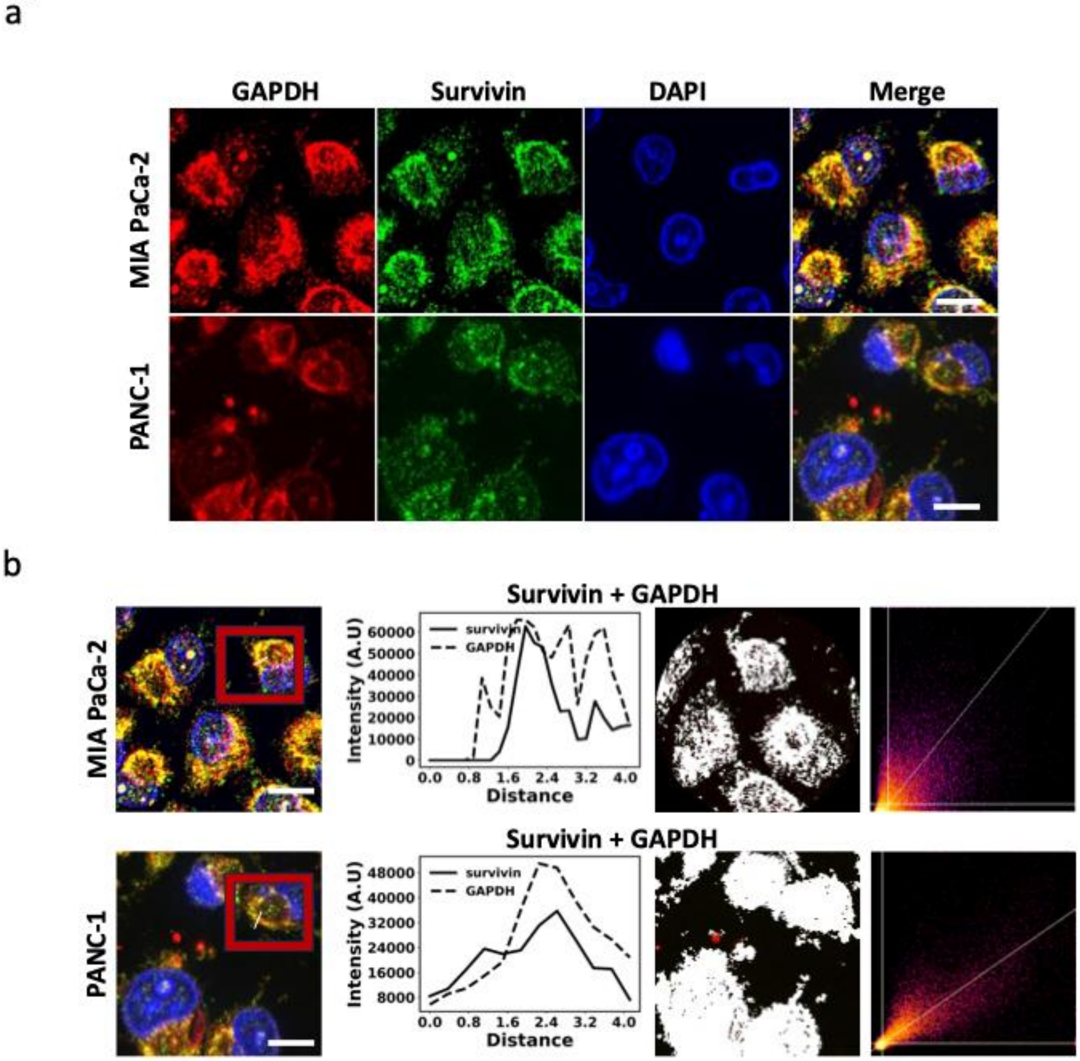


**Figure S6.** a. Full-sized fluorescent microscopy images of MIA PaCa-2 and PANC-1 cells immuno-stained for Survivin and GAPDH. The cells were also stained with DAPI to label the nuclei. These images were used to generate the images of cells shown in Fig. 6a. b. The full co-localization analyses performed on the merged images of the cells shown in Figs. 4a and 4b using ImageJ. The boxed cell shown in each of the first panels was analyzed for the co-localization of the two indicated proteins, with the white line indicating where the analysis was performed. The second panel in each set are graphs showing the extent of co-localization of the two proteins, while the third and fourth panels show the colocalization mask and the colocalization analysis, respectively.
